# *Ehrlichia* SLiM ligand mimetic activates Notch signaling in human monocytes

**DOI:** 10.1101/2022.01.13.476283

**Authors:** LaNisha L. Patterson, Thangam Sudha Velayutham, Caitlan D. Byerly, Duc Cuong Bui, Jignesh Patel, Veljko Veljkovic, Slobodan Paessler, Jere W. McBride

## Abstract

*Ehrlichia chaffeensis* evades innate host defenses by reprogramming the mononuclear phagocyte through mechanisms that involve exploitation of multiple evolutionarily conserved cellular signaling pathways including Notch. This immune evasion strategy is directed in part by tandem repeat protein (TRP) effectors. Specifically, the TRP120 effector activates and regulates Notch signaling through interactions with the Notch receptor and the negative regulator, F-Box and WD repeat domain-containing 7 (FBW7). However, the specific molecular interactions and motifs required for *E. chaffeensis* TRP120-Notch receptor interaction and activation have not been defined. To investigate the molecular basis of TRP120 Notch activation, we compared TRP120 with endogenous canonical/non-canonical Notch ligands and identified a short region of sequence homology within the tandem repeat (TR) domain. TRP120 was predicted to share biological function with Notch ligands, and a function-associated sequence in the TR domain was identified. To investigate TRP120-Notch receptor interactions, colocalization between TRP120 and endogenous Notch-1 was observed. Moreover, direct interactions between full length TRP120, the TRP120 TR domain containing the putative Notch ligand sequence, and the Notch receptor LBR were demonstrated. To molecularly define the TRP120 Notch activation motif, peptide mapping was used to identify an 11-amino acid short linear motif (SLiM) located within the TRP120 TR that activated Notch signaling and downstream gene expression. Peptide mutants of the Notch SLiM or anti-Notch SLiM antibody reduced or eliminated Notch activation and NICD nuclear translocation. This investigation reveals a novel molecularly defined pathogen encoded Notch SLiM mimetic that activates Notch signaling consistent with endogenous ligands.

**Importance:** *E. chaffeensis* infects and replicates in mononuclear phagocytes, but how it evades innate immune defenses of this indispensable primary innate immune cell is not well understood. This investigation reveals the molecular details of a ligand mimicry cellular reprogramming strategy that involves a short linear motif (SLiM) which enables *E. chaffeensis* to exploit host cell signaling to establish and maintain infection. *E. chaffeensis* TRP120 is a moonlighting effector that has been associated with cellular activation and other functions including ubiquitin ligase activity. Herein, we identify and demonstrate that a SLiM present within each tandem repeat of TRP120 activates Notch signaling. Notch is an evolutionarily conserved signaling pathway responsible for many cell functions including cell fate, development, and innate immunity. The proposed study is significant because it reveals the first molecularly defined pathogen encoded SLiM that appears to have evolved *de novo* to mimic endogenous Notch ligands. Understanding Notch activation during *E. chaffeensis* infection provides a model in which to study pathogen exploitation of signaling pathways and will be useful in developing molecularly-targeted countermeasures for inhibiting infection by a multitude of disease-causing pathogens that exploit cell signaling through molecular mimicry.

**Author Summary:** *E. chaffeensis* is a small, obligately intracellular, Gram-negative bacterium that has evolved cellular reprogramming strategies to subvert innate defenses of the mononuclear phagocyte. Ehrlichial TRP effectors interface with the host cell and are involved in pathogen-host interplay that facilitates exploitation and manipulation of cellular signaling pathways; however, the molecular interactions and functional outcomes are not well understood. This study provides molecular insight into a eukaryotic mimicry strategy whereby secreted effectors of obligately intracellular pathogens activate the evolutionarily conserved Notch signaling pathway through a short linear motif ligand mimetic to promote intracellular infection and survival.

## Introduction

*Ehrlichia chaffeensis* is a small, obligately intracellular, Gram-negative tick transmitted bacterium (1) that exhibits tropism for mononuclear phagocytes. *E. chaffeensis* establishes infection through a multitude of cellular reprogramming strategies that involve effector-host interactions resulting the in activation and manipulation of cell signaling pathways to suppress and evade innate immune mechanisms (2–8). The mechanisms whereby *E. chaffeensis* evades host defenses of the macrophage involves exploitation of Wnt and Notch signaling by the tandem repeat protein (TRP) effector, TRP120 (2–6).

*E. chaffeensis* TRP120 effector has well-documented moonlighting functions that include roles as a nucleomodulin (9, 10), a HECT E3 ubiquitin ligase (2, 7, 11), and as a ligand mimic (3, 5, 6, 12). Previously, we found that TRP120 is involved in a diverse array of host cell interactions including components of signaling and transcriptional regulation associated with Wnt and Notch signaling pathways (8). We have recently shown that TRP120 ubiquitinates the Notch negative regulator FBW7 resulting in increased NICD levels, as well as other FBW7 regulated oncoproteins during infection (2). In addition, we have also demonstrated that *E. chaffeensis* Notch activation results in downregulation of toll-like receptor 2 and 4 expression, likely as an immune evasion mechanism (6). Although we have demonstrated TRP120 activates Notch signaling, the molecular details involved in activation have yet to be defined.

The Notch signaling pathway is evolutionarily conserved and is known to play a critical role in cell proliferation, differentiation, and apoptosis in all metazoan organisms (13–16). Notch activation plays significant roles in various other cellular outcomes, including MHC Class II expansion (17), B- and T-cell development (18), and innate immune mechanisms such as autophagy (19) and apoptosis (20, 21). Canonical Notch activation is driven by direct cell-membrane bound receptor-ligand interactions with four Notch receptors (Notch1-4) and canonical Notch ligands, Delta-like (DLL 1,3,4) and Jagged (Jagged/Serrate-1 and 2). Notch receptor-ligand interactions occur at the Notch extracellular domain (NECD), specifically at epidermal growth factor-like repeats (EGFs) 11-13, the known ligand binding domain (LBD). Module at the N-terminus of Notch ligands (MNNL) and Delta/Serrate/LAG-2 (DSL) domains in canonical Notch ligands interact with the Notch LBD. Although there is evidence demonstrating the requirement of both N-terminal MNNL and DSL Notch ligand domains for Notch receptor binding, there is little information known about ligand regions/motifs that are necessary for Notch activation (22, 23). During canonical Notch activation, ligands expressed on neighboring cells bind the Notch receptor and create a mechanical force at the negative regulatory region (NRR) which triggers several sequential proteolytic cleavages, releasing the Notch intracellular domain (NICD). NICD subsequently translocates to the nucleus and binds to other transcriptional coactivators, including RBPjK and MAML, to activate Notch gene transcription. Notably, secreted non-canonical Notch ligands have also been shown to activate Notch signaling; however, the molecular details of non-canonical Notch ligand-receptor interactions are not well defined.

There are three major classes of protein interaction modules which include globular domains, IDDs, and short linear motifs (SLiMs), all of which have distinct biophysical attributes (24–26). IDDs are 20-50 amino acids in length, are known to be disordered in nature, are located within globular domains or intrinsically disordered protein regions and have transient interactions in the nanomolar range. In comparison, SLiMs are ~3-12 amino acids in length, are known to be disordered in nature, located within globular domains or IDDs, and have low micromolar affinity ranges with transient interactions. SLiMs have been shown to evolve *de novo* for promiscuous binding to various partners (26, 27). Ehrlichial TRPs interact with a diverse array of host proteins through several well-known protein-protein interaction mechanisms including post-translational modifications (PTMs), and various protein interaction modules located in intrinsically disordered domains (IDDs) (7, 9, 26, 28).

Microorganisms have developed mechanisms to survive in the host cell which involve hijacking host cell processes. Molecular mimicry has been well-established as an evolutionary survival strategy utilized by pathogens to disrupt or co-opt host function for infection and survival [26-29]. Studies have determined this occurs through pathogen effectors that mimic eukaryotic host proteins, allowing for pathogens to hijack and manipulate host cellular pathways and functions. SLiMs have been identified as interaction modules whereby eukaryotes and pathogens direct cellular processes through protein-protein interactions (29, 30). Recently, we have demonstrated TRP120 is a Wnt ligand mimetic that interacts with host Wnt receptors to activate Wnt signaling (3).

In this study, we reveal an *E. chaffeensis* Notch SLiM ligand mimetic whereby TRP120 activates Notch signaling for infection and intracellular survival. Understanding the molecular mechanisms utilized by *E. chaffeensis* to subvert innate host defense for infection and survival is essential for understanding intracellular pathogen infection strategies and provides a model to investigate molecular host-pathogen interactions involved in repurposing host signaling pathways for infection.

## Results

### *E. chaffeensis* TRP120 shares sequence homology and predicted Notch ligand function

We have previously shown TRP120 interacts with Notch activating metalloprotease, ADAM17 and Notch antagonist FBW7 using yeast-two hybrid analysis (Y2H) (8). We have also shown that TRP120 binds to the promoter region of *notch1* using chromatin immunoprecipitation sequencing (ChIP-Seq), and that activation of Notch occurs during infection (6, 10). Notch activation occurs through direct interaction of Notch ligands with the Notch-1 receptor initiating two receptor proteolytic cleavages, resulting in NICD nuclear translocation and subsequent activation of Notch downstream targets. Since TRP120 has been shown to activate the Notch signaling pathway, we examined TRP120 sequence homology and correlates of biological functionality with Notch ligands.

NCBI Protein Basic Local Alignment Search Tool (BLAST) was used to identify local similarity between TRP120 and canonical/non-canonical Notch ligand sequences. Sequence homology with a TRP120 tandem repeat (TR) IDD motif, TESHQKEDEIVSQPSSE (aa. 284-301), was shown to share sequence homology with several canonical Notch ligands, including Jagged-1, DLL1, DLL4, and non-canonical Notch ligand TSP2 (Fig. 1A). We then used informational spectrum method (ISM) to predict similar functional properties between TRP120 and Notch ligands. ISM is a prediction method that uses the electron ion interaction potential of each amino acid within the primary sequence of proteins to translate the primary sequences into numerical sequences. Translated sequences are then converted into a spectrum using Fourier transform. Cross spectral analysis of the translated sequences is then performed to obtain characteristic frequency peaks that demonstrate if proteins share a similar biological function. TRP120 was predicted to share a similar biological function with canonical Notch ligands, DLL1, 3 and 4, and non-canonical Notch ligand F3 contactin-1, a known adhesion molecule (Figs. S1 A-D). To identify the sequence responsible for the identified frequency peaks, reverse Fourier transform of ISM was performed (Fig. 1B). A 35-mer TRP120-TR IDD motif, IVSQPSSEPFVAESEVSKVEQEETNPEVLIKDLQD (aa. 214-248 and 294-328), was associated with characteristic frequency peaks (Figs. 1B and 1C). Collectively, these results indicate that the TRP120 sequence and fundamental biophysical properties of the amino acids are consistent with Notch ligands.

**Fig. 1.**
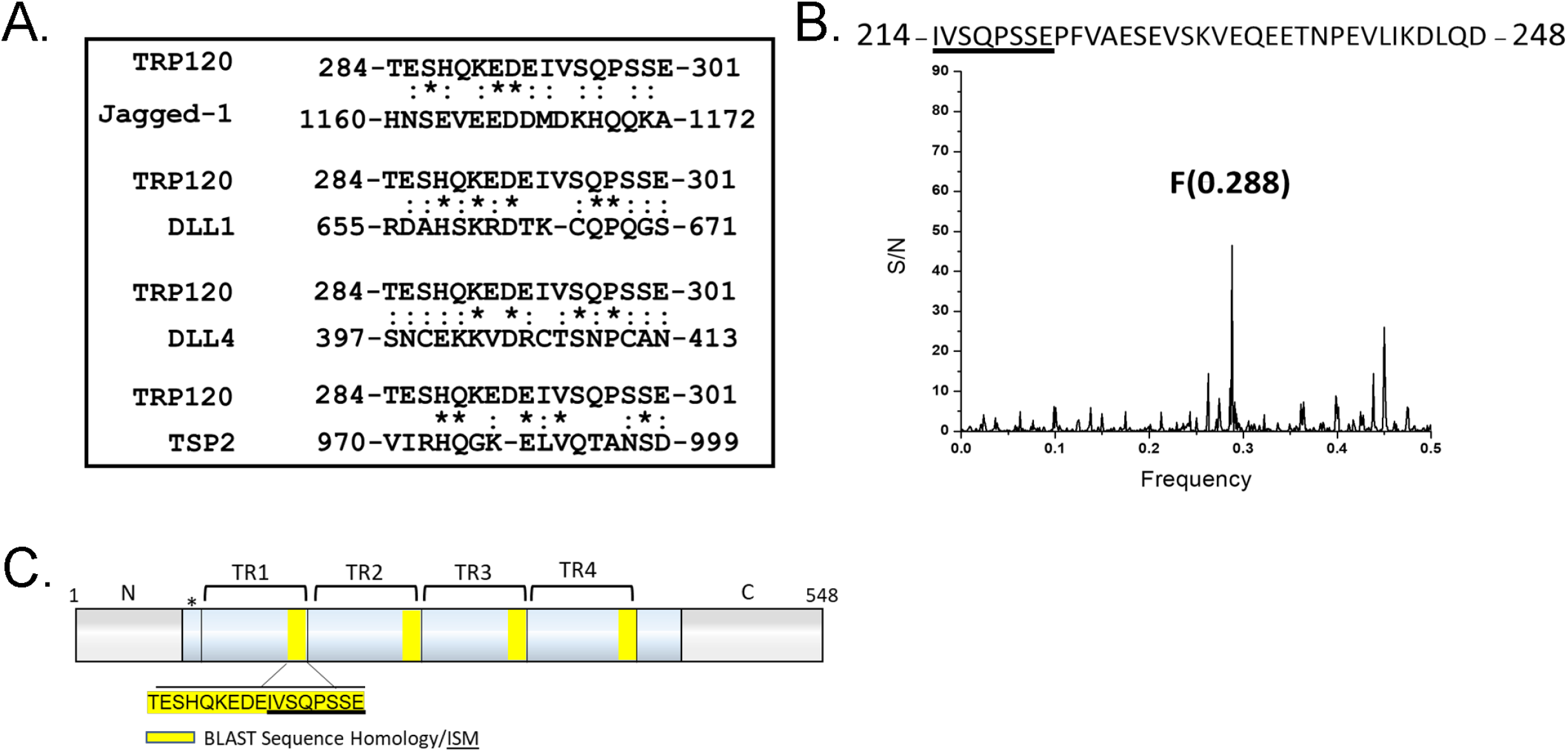
*E. chaffeensis* TRP120 shares sequence homology and biological function with canonical and noncanonical Notch ligands. (A) BLAST analysis of TRP120 with canonical/noncanonical Notch ligands demonstrating amino acid homology. An asterisk (*) represents identical conserved amino acid residues; a colon (:) represents conservative substitutions. (B) Informational Spectrum Method (ISM) was used to predict if TRP120 shared similar biological function with canonical and nonocanonical Notch ligands. Primary sequences of TRP120 and Notch ligands were converted into a numerical sequence-based electron ion interaction potential (EIIP) of each amino acid. Numerical sequences were converted into a spectrum using Fourier transform. To determine if proteins shared a similar biological function and cross spectra analysis was performed and similar biological function is denoted by a peak at a frequency of F(0.288). (C) Schematic of TRP120 N- C- (gray) and TR domains (blue) with four highlighted repetitive TRP120 TR motifs that share sequence homology with Notch ligands. ISM sequence shown in panel B (underlined). (*) represents a partial tandem repeat containing similar (EDDTVSQPSLE) but non identical sequence to highlighted sequence.

### *E. chaffeensis* TRP120 directly interacts with the Notch-1 ligand binding region (LBR)

Canonical activation of the Notch pathway is known to occur through canonical Notch ligands binding to Notch receptor LBD (EGFs 11-13 in the extracellular domain). To investigate if TRP120 interacts with the Notch-1 receptor LBR (EGFs 1-15), we ectopically expressed GFP-tagged full length TRP120 (TRP120-FL-GFP) in HeLa cells and probed for endogenous Notch-1 to determine colocalization. Pearson’s correlation coefficient (PC) and Mander’s coefficient (MC) (correlation range +1 to −1; 0 represents absence of correlation), was used to quantify the degree of colocalization between TRP120-FL-GFP and Notch-1. Ectopically expressed TRP120-FL-GFP was found to strongly colocalize (PC = 0.897 and MC = 0.953) with endogenous Notch-1 (Fig. 2A). Colocalization of TRP120 and Notch-1 demonstrates that these two proteins are in the same spatial location; however, it does not demonstrate direct protein-protein interaction. To confirm a direct interaction, we utilized pull-down assays of TRP120-FL and Notch-1 LBR. A His-tagged rTRP120-FL (rTRP120-FL-His) construct was incubated with a Fc-tagged recombinant Notch-1 LBR, and a direct protein-protein interaction was demonstrated (Figs. 2B and S2A-B). Thioredoxin (TRX), used as a fusion tag in the pBAD expression vector containing TRP120 constructs and as a recombinant control, did not interact (Fig. 2B). Based on sequence homology and ISM data, a short region of sequence homology within the tandem repeat (TR) domain was identified that could be involved in the TRP120 and Notch-1 LBR. To determine if the TRP120-TR was responsible for the previous TRP120 and Notch-1 LBR interaction, we performed a pull-down assay with TRP120-TR and Notch-1 LBR. rTRP120-TR was pulled down with anti-Notch-1 LBR antibody demonstrating a direct interaction with the TR domain (Fig. 2C).

**Fig. 2.**
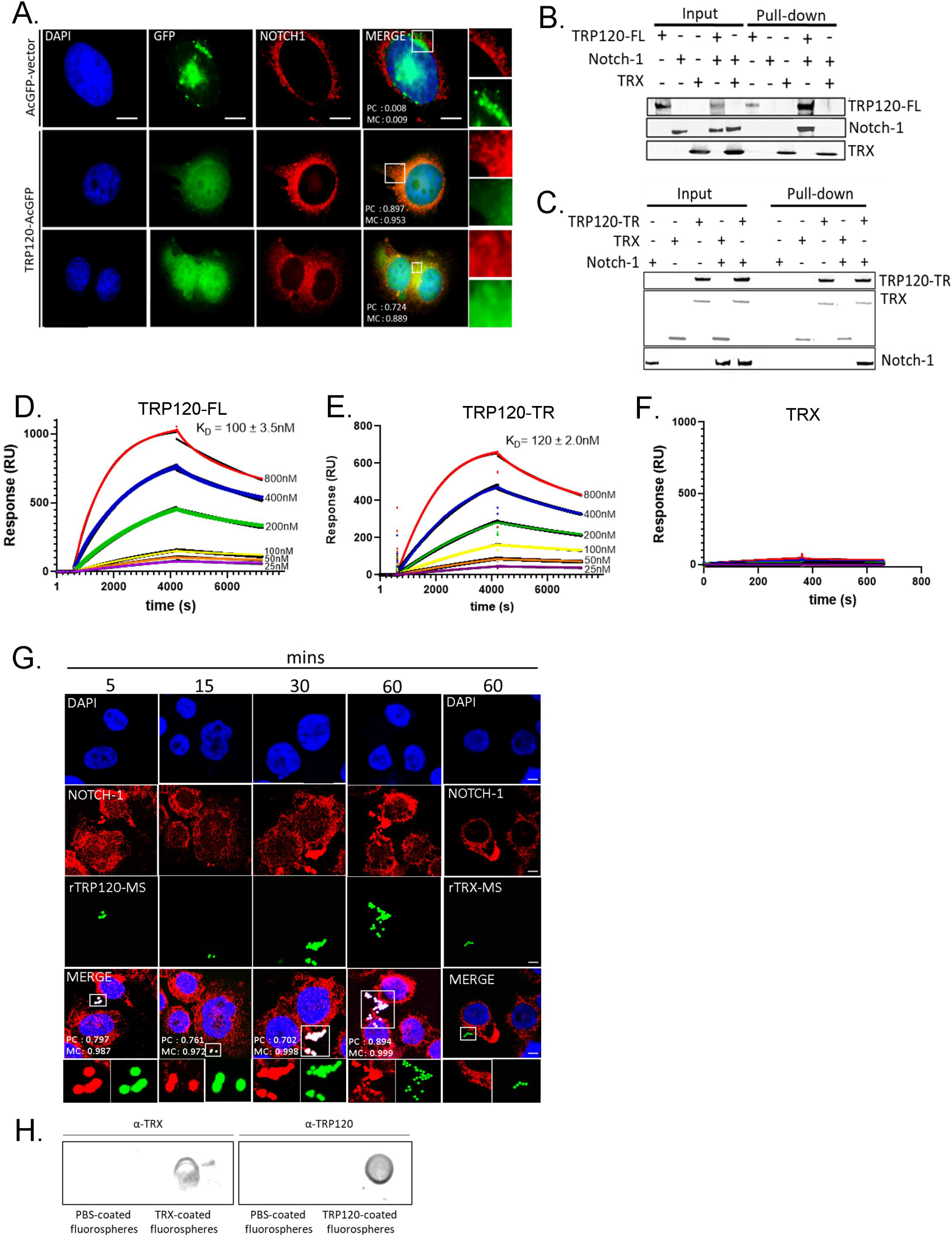
*E. chaffeensis* TRP120-TR interactions with the Notch receptor ligand binding region (LBR). (A) HeLa cells transfected with TRP120-GFP (green) and probed for endogenous Notch-1 (red) demonstrate colocalization by immunofluorescent microscopy. Colocalization was quantitated by Pearson’s and Mander’s coefficient (−1 no colocalization; +1 strong colocalization). (B and C) His-tag pull down assays demonstrating direct interaction between TRP120 and Notch-1. Recombinant Fc-tagged Notch-1 LBR was incubated with (B) TRP120-FL-His, (C) TRP120-TR-His or TRX-His negative control on Talon metal affinity resin. Bound Notch-1, TRP120-His, α-TRP120 against a TR peptide or TRX-His were detected with α-Notch-1, α-TRP120 or α-TRX antibodies. (D-F) Surface plasmon resonance of (D) TRP120-FL-His, (E) TRP120-TR-His or (F) TRX-His with Fc-tagged Notch-1 LBR on a Biacore T100 with a series S Ni-nitrilotriacetic acid (NTA) sensor chip. TRP120-FL-His, TRP120-TR-His or TRX-His were immobilized on the NTA chip and 2-fold dilutions (800nM to 25nM) of Fc-tagged Notch-1 LBR were used as analyte to determine binding affinity (K_D_). Sensograms and K_D_ are representative of data from triplicate experiments. (G) THP-1 cells were treated with rTRX- or rTRP120-FL-coated fluorescent microspheres for varying time points (5-60 mins). Colocalization was visualized by confocal immunofluorescent microscopy. Notch-1 was immunostained with tetramethylrhodamine isothiocyanate [TRITC] and TRP120-coated fluorescein isothiocyanate [FITC] auto-fluorescent microspheres. Nuclei were stained with DAPI (blue). White boxes indicate areas of colocalization measurements. Scale bar = 10 μm. (H) Dot blot of PBS, TRX or TRP120-FL-coated microspheres probed with α-TRX or α-TRP120 antibodies, respectively.

To further confirm direct interaction of TRP120-FL or TRP120-TR and Notch-1 LBR, surface plasmon resonance was performed. An interaction between both rTRP120-FL (Fig. 2D) and rTRP120-TR (Fig. 2E) with Notch-1 LBR was detected in a concentration dependent manner. Fitting the concentration response plots for TRP120-FL and TRP120-TR yielded a K_D_ (equilibrium dissociation constant) of 100 ± 3.5 nM and 120 ± 2.0 nM, respectively (Figs. 2D-E). No interaction was detected between TRX and Notch-1 LBR (Fig. 2F). Additionally, treatment of THP-1 cells with TRP120-coated sulfate, yellow-green microspheres demonstrated colocalization of TRP120 and Notch-1 (Figs. 2G-H). In comparison, TRX-coated fluorescent microsphere did not colocalize with the Notch-1 receptor (Figs. 2G-H). Together, these binding data reveal TRP120-TR binds the Notch-1 LBR.

### *E. chaffeensis* TRP120-TR domain is required for Notch activation

Both the N-terminal MNNL and cysteine-rich DSL domain of Notch ligands are known to be required for receptor binding; however, there is little known regarding ligand motifs required for Notch activation. We have previously demonstrated Notch activation occurs in THP-1 cells after stimulation with TRP120-coated beads for 15 min (6). Gene expression levels of *notch1, hes1* and *hes5* were upregulated after incubation with TRP120-coated beads. To further delineate the TRP120 domain required for Notch activation THP-1 cells or primary human monocytes were treated with soluble purified full length or truncated constructs of recombinant TRP120 (rTRP120-TR and -C-terminus) (Figs. S2A-B). Full length rTRP120 and rTRP120-TR caused NICD nuclear translocation 2 h post-treatment (Figs. 3A-B). NICD nuclear translocation was not observed in untreated cells, cells treated with TRX or rTRP120-C-terminal soluble proteins (Figs. 3A-B). Collectively, these data demonstrate the requirement of TRP120-TR for Notch activation.

**Fig. 3.**
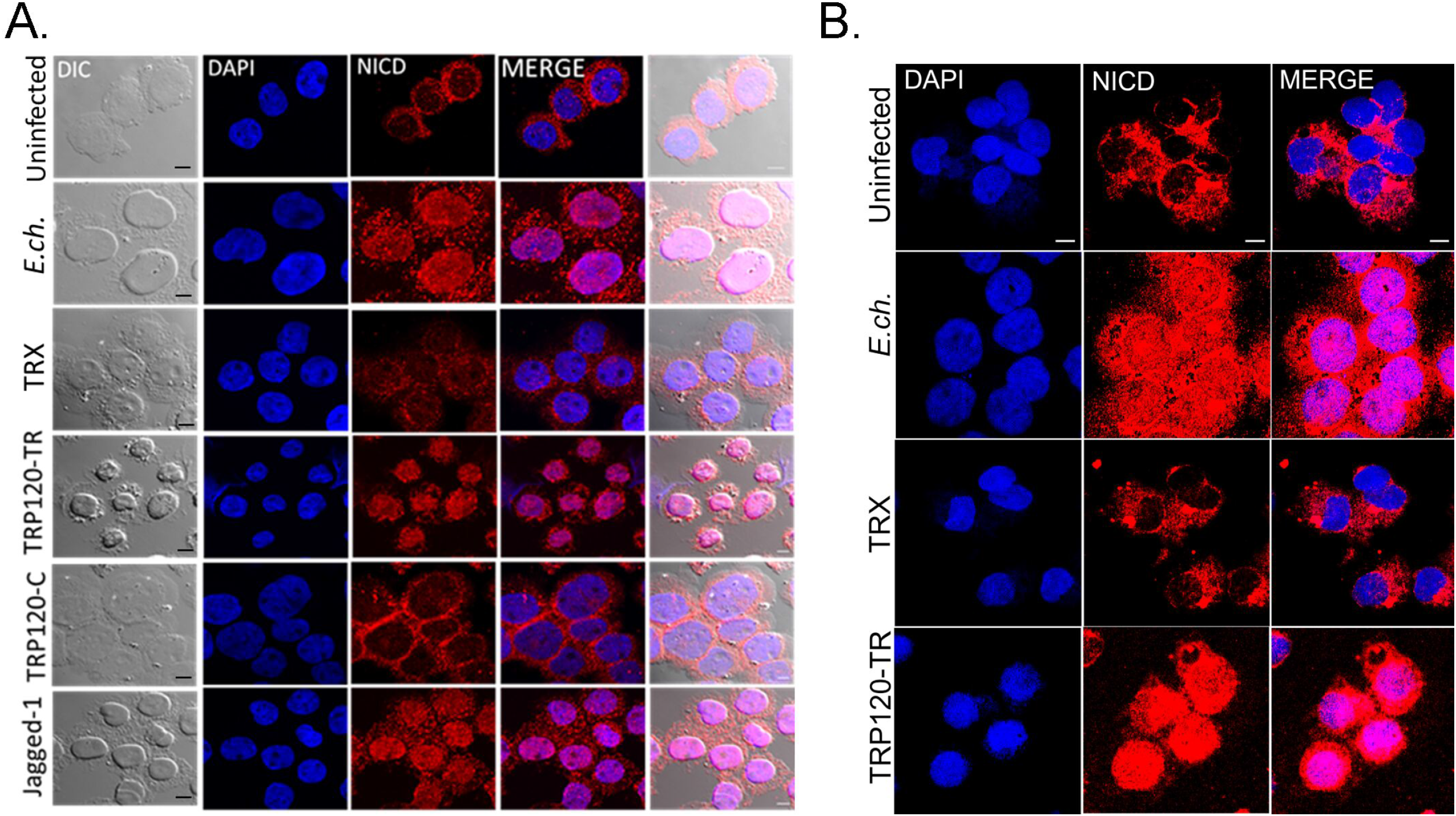
TRP120-TR activates Notch and NICD nuclear translocation in primary human monocytes. (A) Soluble recombinant TRP120-TR or -C terminal proteins (2 μg/ml) were incubated with THP-1 cells for 2 h. Cells were collected and NICD localization determined by confocal immunofluorescent microscopy. Uninfected/untreated or recombinant TRX-treated THP-1 cells were used as negative controls. *E. chaffeensis*-infected or recombinant Jagged-1 treated THP-1 cells were used as positive controls. NICD nuclear translocation was detected in *E. ch*-infected, TRP120-TR and Jagged-1 treated cells. (B) Primary human monocytes were treated with soluble TRP120-TR or recombinant TRX as described above and NICD nuclear translocation was detected in *E. chaffeensis-infected* and TRP120-TR-treated cells. End point analysis was performed as described in Fig. 2. Experiments were performed in triplicate and representative images are shown.

### *E. chaffeensis* TRP120-TR Notch ligand IDD-mimetic activates Notch

To determine if Notch is activated by a TRP120-TR Notch mimetic IDD motif, several TRP120-TR synthetic peptides were generated (Fig. 4A). THP-1 cells or primary human monocytes were treated with TRP120-TR IDD peptides or scrambled negative control peptide for 2 h. A 35-aa TRP120-TR IDD motif (TRP120-N1-P3) caused nuclear translocation (Figs. 4B and C). Importantly, the identified IDD contained a motif identified in both sequence homology and ISM data (Fig. 1C). Inhibition of Notch signaling by DAPT, a γ-secretase inhibitor, abrogated Notch activation with TRP120-N1-P3 treatment, indicating that TRP120-N1-P3 directly binds to the Notch-1 receptor for Notch activation (Fig. 4B). To confirm Notch activation by TRP120-N1-P3, gene expression levels of Notch downstream targets were examined by human Notch signaling pathway array analysis. In comparison to untreated THP-1 cells, a significant increase in Notch downstream targets, including HES1, HES5, HEY1 and HEY2 gene expression levels occurred in TRP120-N1-P3 treated cells (Figs. 5A-B, Table S1A). Interestingly, Notch gene expression by TRP120-N1-P3 treatment was increased in a concentration-dependent manner (Fig. 5B, Table S1A). Importantly, rJagged-1 also demonstrated similar upregulation of Notch genes in a concentration-dependent manner (Fig. S3). These data demonstrate that a TRP120 IDD mimetic motif is responsible for TRP120 Notch activation.

**Fig. 4.**
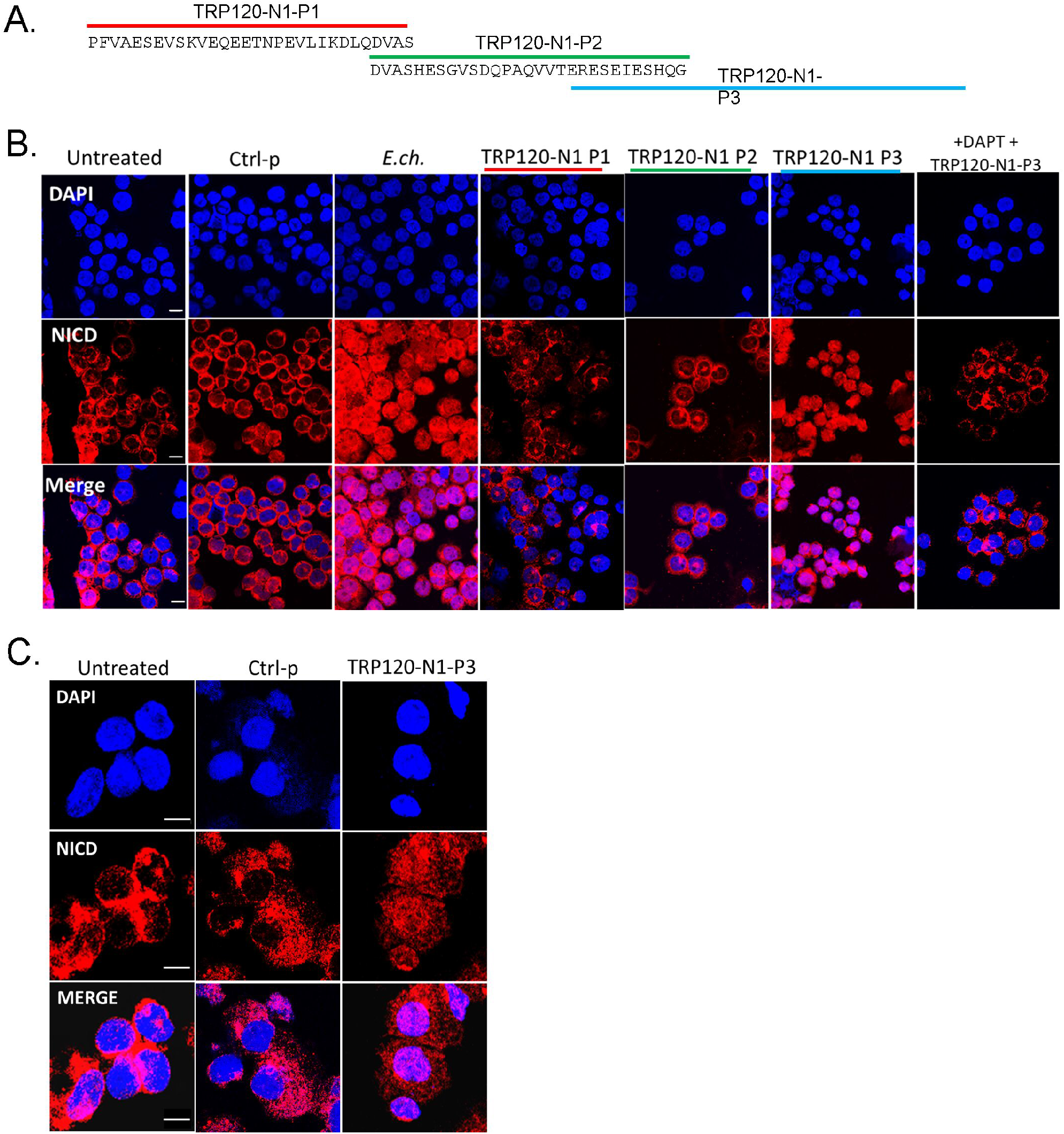
A TRP120-TR Notch-1 memetic IDD peptide stimulates NICD nuclear translocation. (A) Overlapping TRP120-TR IDD peptide sequences (P1-P3) (B) THP-1 cells or (C) Primary human monocytes were incubated with synthetic TRP120-TR IDD peptides to determine the TRP120-TR Notch-1 memetic motif responsible for Notch activation. TRP120-TR peptides were overlapping peptides spanning an entire TR domain. Cells were treated with peptide (1 μg/ml) for 2 h and confocal immunofluorescent microscopy was used to visualize NICD localization. NICD nuclear translocation denotes Notch activation. A scrambled peptide (Ctrl-p) was used as negative control and *E.ch*. infected cells were used as positive control. To determine if direct interaction of the TRP120-N1-P3 peptide and Notch receptor was necessary for Notch activation, THP-1 cells were pre-treated with DAPT, a γ-secretase inhibitor, and treated with TRP120-N1-P3 peptide for 2 h.

**Fig. 5.**
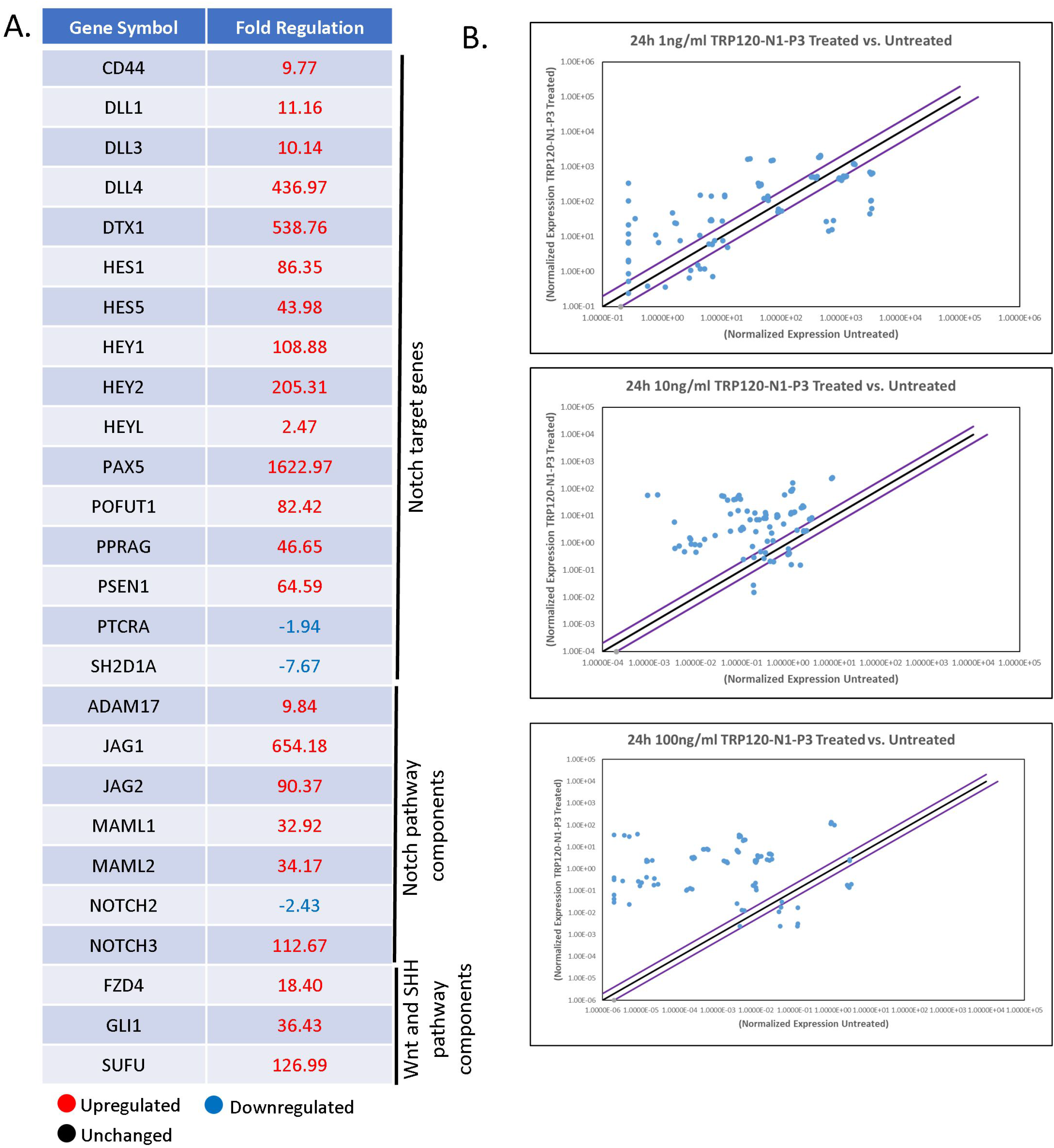
TRP120-N1-P3 IDD peptide stimulates Notch gene expression. (A) Table of Notch pathway genes with corresponding fold-change displaying differential expression (up, down or no change) at 24 h p.t with 10 ng/ml of TRP120-N1-P3 peptide (B) Scatter plots of expression array analysis of 84 Notch signaling pathway genes to determine Notch gene expression 24 h after stimulation with 1 ng/ml (top), 10 ng/ml (middle) or 100 ng/ml (bottom) of TRP120-N1-P3 peptide. Purple lines denote a 2-fold up or down regulation in comparison to control, and the black line denotes no change. Scatter plots are representative of three independent experiments (n = 3).

### *E. chaffeensis* TRP120-TR Notch ligand SLiM mimetic activates Notch

It is well-documented that SLiMs are found in two general groups; posttranslational modification (PTM) motifs or ligand motifs that mediate binding events. We have previously identified a functional TRP120 HECT E3 ligase catalytic motif located in the C-terminus (2, 7) and have recently identified a TRP120-TR Wnt SLiM mimetic motif (3). To determine if the TRP120-TR Notch mimetic motif could be a SLiM (3-12 aa), overlapping TRP120-TR synthetic peptides that span the identified 35-aa TRP120-TR IDD motif were synthesized (Fig. 6A). Treatment with P4 and P5 TRP120-TR Notch mimetic SLiM peptides in THP-1 cells did not result in NICD nuclear translocation (Fig. 6B); however, TRP120-TR Notch mimetic SLiM P6 (TRP120-N1-P6) located at the C-terminus resulted in NICD nuclear translocation (Fig. 6B). TRP120-N1-P6 was also shown to cause NICD nuclear translocation in primary human monocytes (Fig. 6C). Furthermore, pre-treatment of DAPT inhibited TRP120-N1-P6 Notch activation (Fig. 6B). Upregulation of Notch downstream targets occurred with TRP120-N1-P6 treatment in a concentration dependent manner (Figs. 7A-B, Table S1B), as previously shown with the TRP120-N1-P3 peptide. In comparison, TRP120-N1-P5 peptide treatment, did not result in significant upregulation of Notch gene expression (Fig. 7B).

**Fig. 6.**
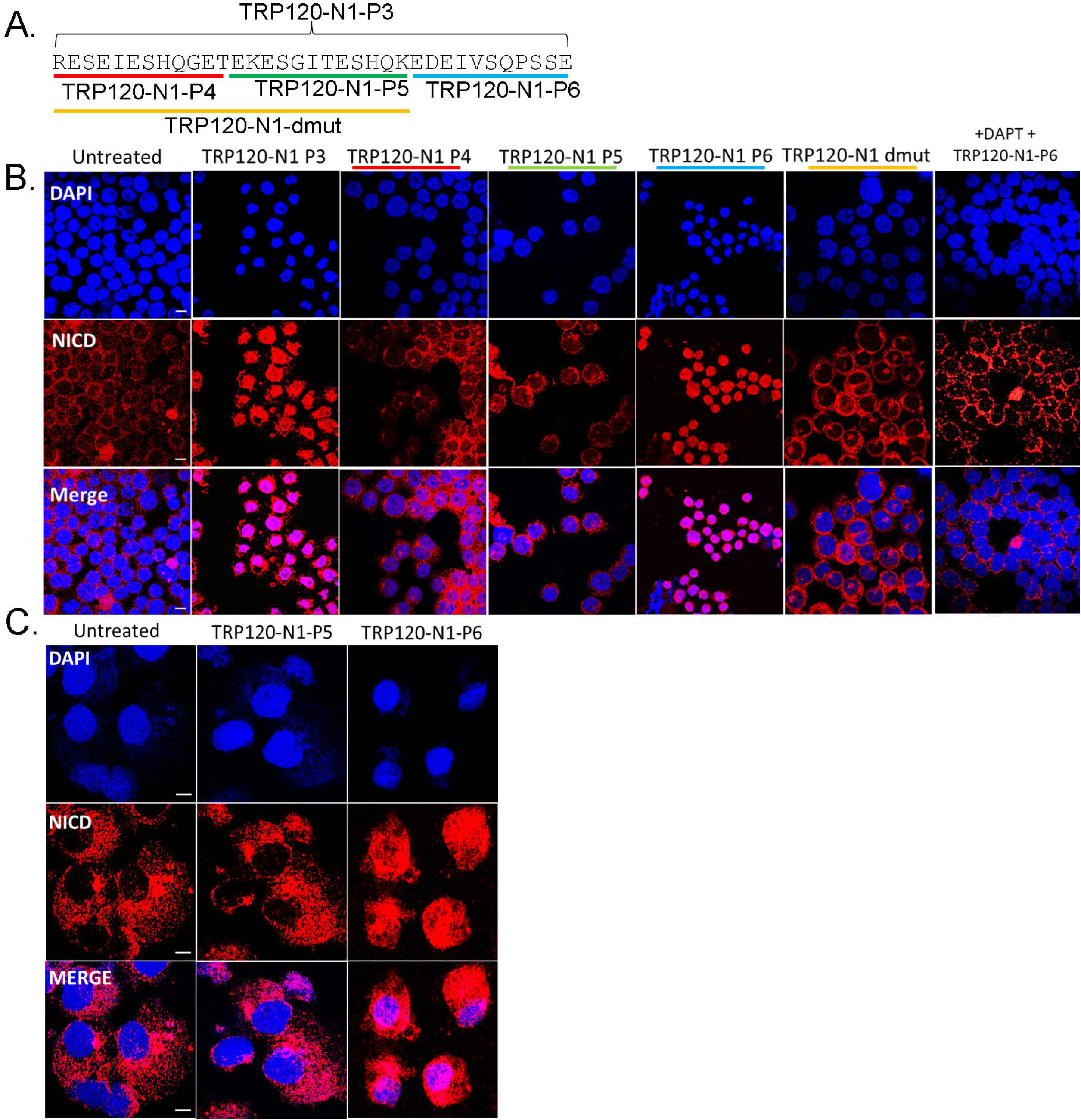
A TRP120-TR Notch-1 memetic SLiM peptide activates Notch signaling. (A) TRP120-N1 SLiM (P4-P6) and mutant (dmut) peptide sequences. (B) THP-1 cells or (C) primary human monocytes were treated with synthetic TRP120-TR SLiM peptides to identify the TRP120-TR Notch-1 SLiM memetic motif. TRP120-TR peptides were SLiM peptides spanning the entire TRP120-N1-P3 peptide sequence. TRP120-N1-P3 mutant peptide (dmut) has a deletion of the TRP120-N1-P6 amino acids. Cells were treated with peptide (1 μg/ml) for 2 h and NICD localization visualized by confocal microscopy. TRP120-N1-P3 peptide was used as a positive control. To determine if direct interaction of the TRP120-N1-P6 peptide and Notch receptor was necessary for Notch activation, THP-1 cells were pre-treated with DAPT, a γ-secretase inhibitor, and treated with TRP120-N1-P6 peptide for 2 h. Representative data of all experiments are shown (n = 3).

**Fig. 7.**
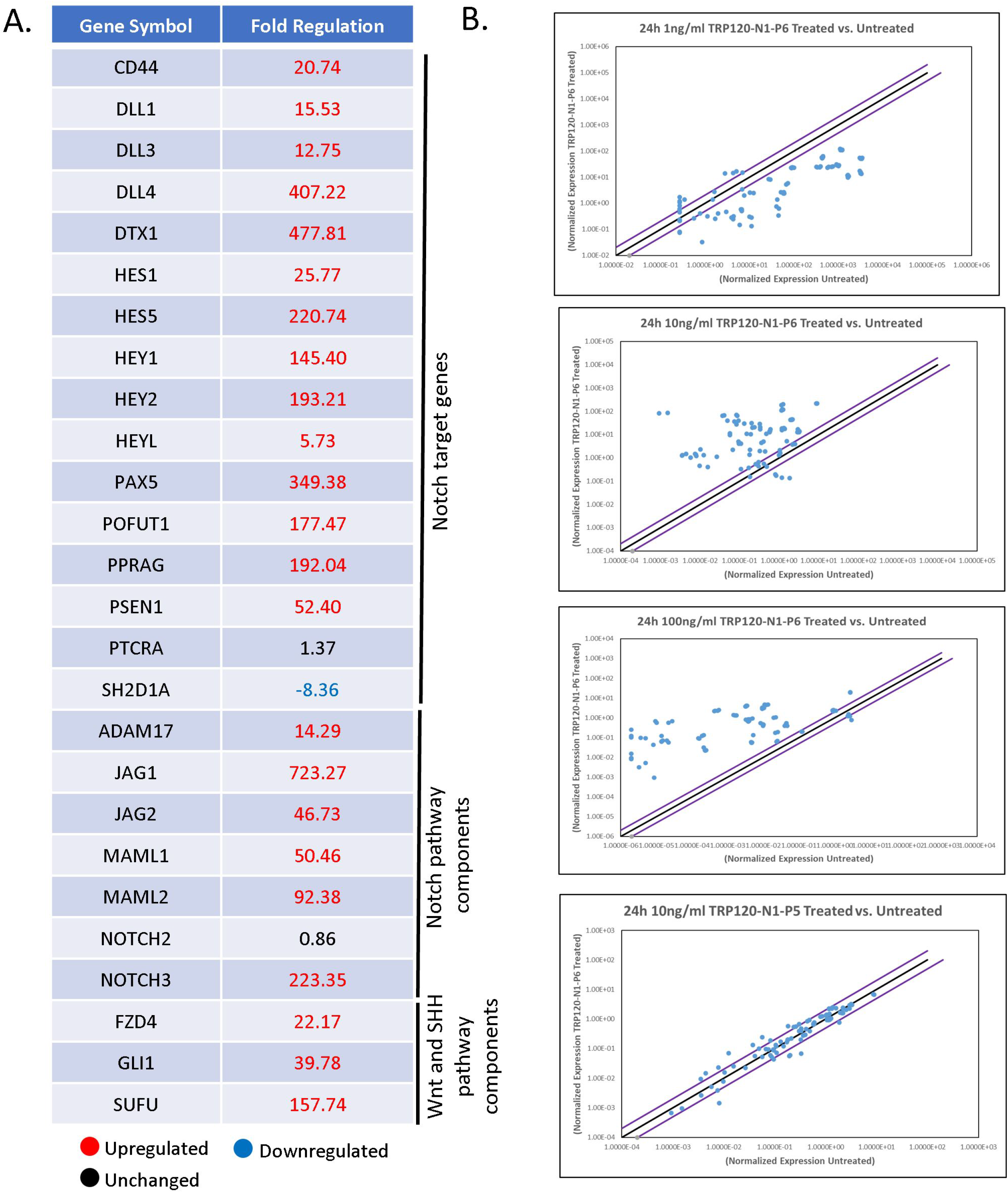
TRP120-N1-P6 SLiM Notch memetic peptide stimulates Notch gene expression. (A) Selected Notch pathway genes with corresponding fold-change displaying differential expression (up- and downregulation) at 24 h p.t with 10 ng/ml of TRP120-N1-P6 peptide. (B) Scatter plots of expression array analysis of 84 Notch signaling pathway genes to determine Notch gene expression with 1 ng/ml, 10 ng/ml, 100 ng/ml of TRP120-N1-P6 peptide or TRP120-N1-P5 treatment (10 ng/ml) compared to untreated cells (bottom) at 24 p.t. Purple lines denote a 2-fold up or down regulation in comparison to control, and the black lines denotes no change. Scatter plots are representative of three independent experiments (n = 3).

To confirm that TRP120-N1-P6 is required for Notch activation, a TRP120-N1-P3 mutant peptide (dmut) (Fig. 6A) without the TRP120-N1-P6 motif was tested. THP-1 cells stimulated with TRP120-N1-dmut exhibited abrogated Notch activation as demonstrated by NICD translocation (Fig. 6B). To determine the minimal residues required in the TRP120-TR Notch mimetic SLiM, alanine mutagenesis was used to determine the contribution of specific residues to Notch activation (Fig. 8A, blue boxes). Mutated residues were selected based on sequence homology and ISM data. Mutants (dmut-1 −2, −3 and −4) exhibited reduced Notch activation as determined by NICD translocation, but only the TRP120-N1-dmut peptide resulted in full abrogation of NICD nuclear translocation (Fig. 8A). Collectively, these data demonstrate that the TRP120-N1-P6 SLiM is a Notch mimetic.

**Fig. 8.**
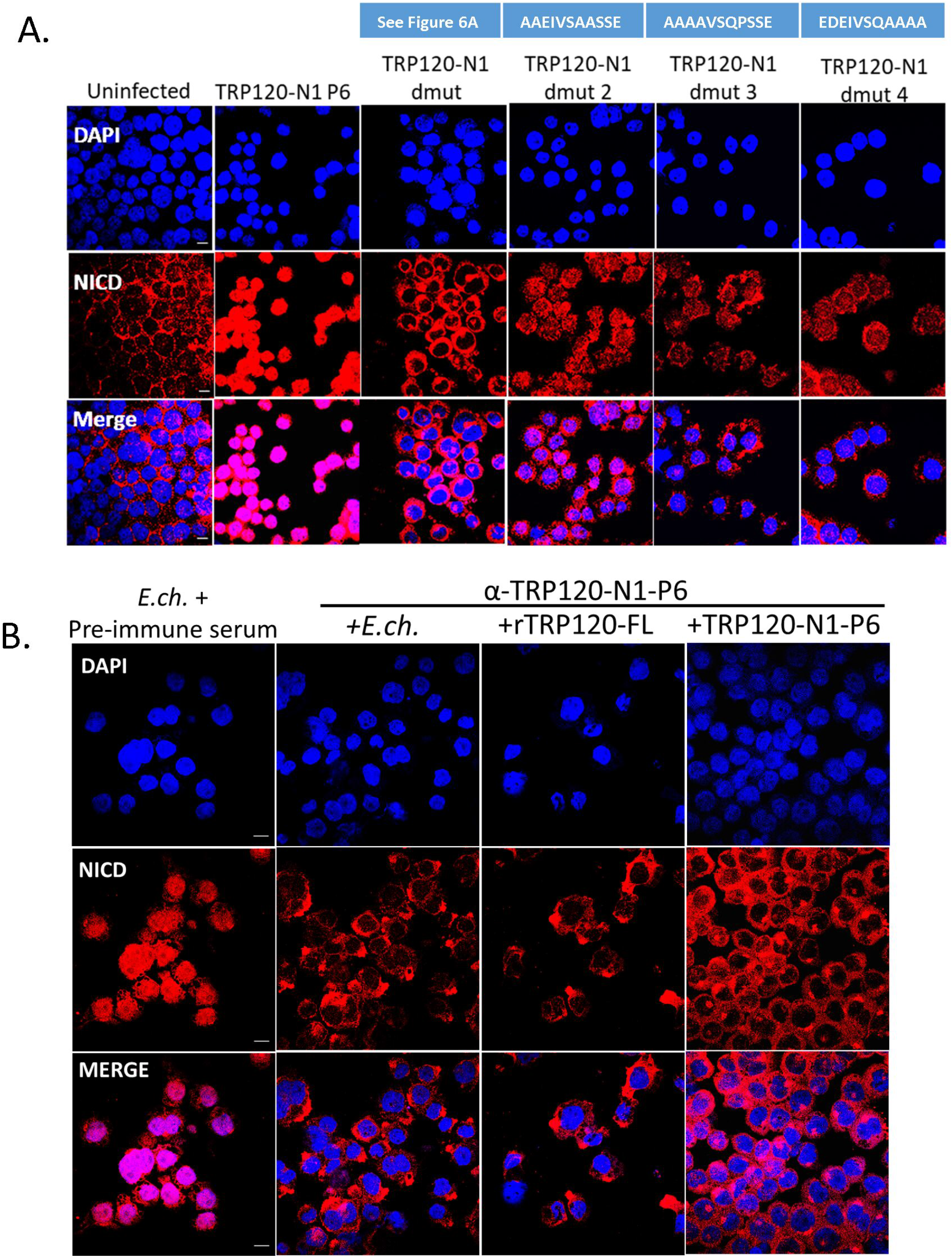
Amino acids critical to TRP120-N1-P6 memetic SLiM activity and anti-SLiM antibody blocks Notch activation. (A) Critical amino acids of the TRP120-N1-P6 memetic SLiM determined by alanine mutagenesis (mutant peptide sequences are shown above the corresponding panel). THP-1 cells were treated with mutant peptides (dmut2, −3 and −4; 1 μg/ml) for 2 h and confocal immunofluorescent microscopy was used to visualize NICD localization. NICD nuclear translocation denotes Notch activation. Peptide dmut was used as a negative control and TRP120-N1-P6 was used as a positive control. (B) Cell-free *E. chaffeensis*, rTRP120-FL or TRP120-N1-P6 were incubated with α-TRP120-N1-P6 rabbit polyclonal antibody (5 μg/ml) for 30 min. Preimmune serum was used as control antibody. THP-1 cells were subsequently inoculated with the cell-free *E. chaffeensis/α-TRP120-N1-P6* mixture for 2 h and confocal immunofluorescent microscopy was used to visualize NICD nuclear localization. Representative data of all experiments are shown (n = 3).

**Fig. 9.**
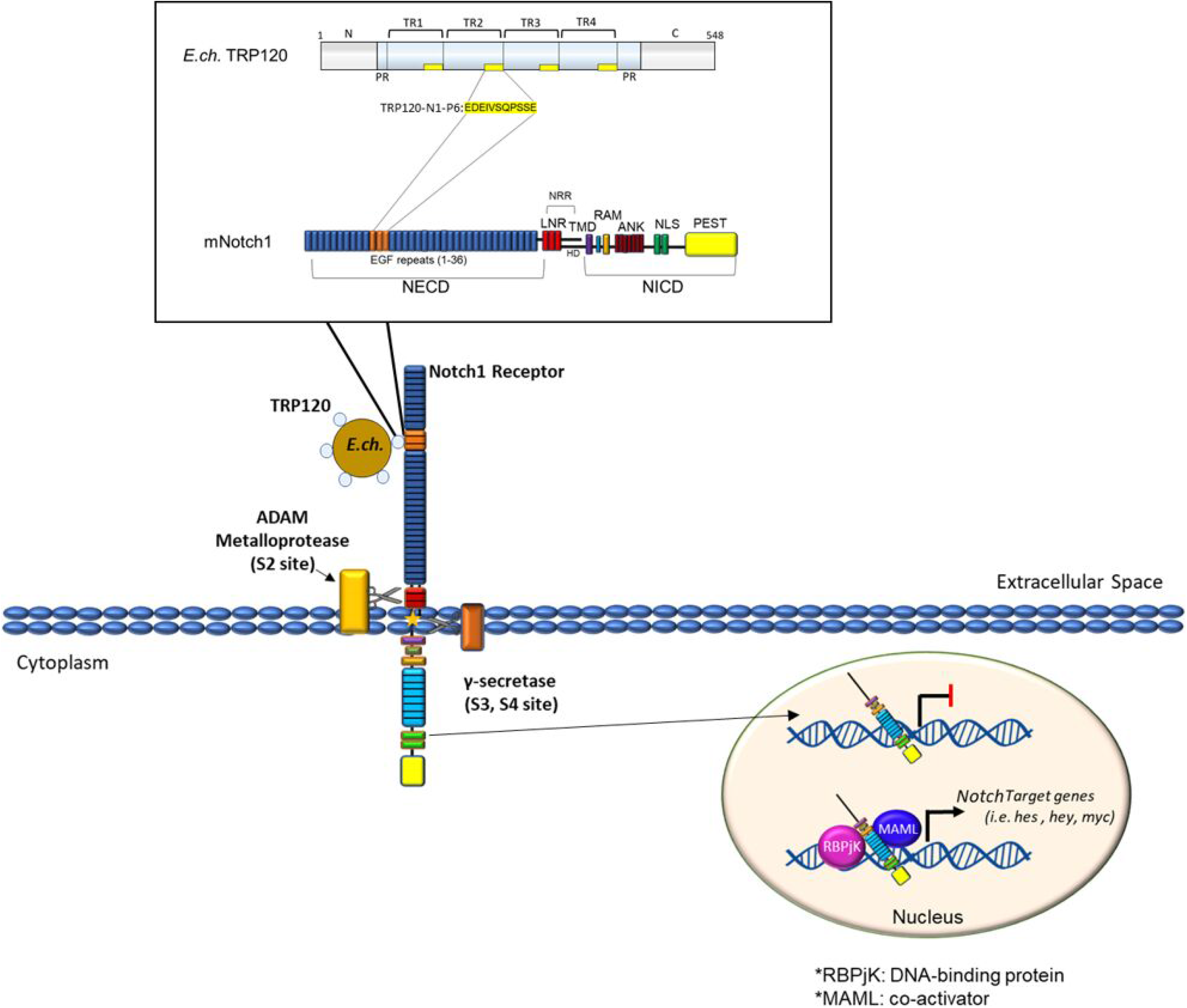
Proposed model of *E. chaffeensis* TRP120 Notch activation. A TRP120-TR Notch SLiM memetic motif (TRP120-N1-P6; yellow highlight) binds the Notch-1 extracellular domain at a region containing the confirmed Notch ligand binding domain (LBD) to activate Notch signaling. TRP120-N1-P6 binding leads to NICD nuclear translocation and upregulation of Notch gene targets.

### TRP120 Notch SLiM antibody blocks *E. chaffeensis* Notch activation

To investigate whether the TRP120 Notch mimetic is solely responsible for Notch activation by *E. chaffeensis*, THP-1 cells were pre-treated with a purified rabbit polyclonal antibody generated against the TRP120-N1-P6 SLiM and subsequently infected with *E. chaffeensis* for 2 h. Negative pre-immune serum was used as a negative control. NICD nuclear translocation was determined in α-TRP120-N1-P6 SLiM or negative pre-immune serum treated cells. THP-1 cells treated with α-TRP120-N1-P6 SLiM did not display NICD nuclear translocation, in comparison to the negative pre-immune serum control (Fig. 8B). These data suggests that TRP120-N1-P6 SLiM is the only Notch mimic involved in Notch activation by *E. chaffeensis*.

## Discussion

We have previously demonstrated TRP120-host interactions to occur with a diverse array of host cell proteins associated with conserved signaling pathways, including Wnt and Notch (8). Two proteins shown to interact with TRP120 were the Notch metalloprotease, a disintegrin and metalloprotease domain (ADAM17), and a Notch antagonist, F-box and WD repeat domain-containing 7 (FBW7). In addition, we have demonstrated that secretion of *E. chaffeensis* TRP120 activates Notch signaling to downregulate TLR2/4 expression for intracellular survival. Moreover, Keewan et al, demonstrated upregulation of Notch-1, IL-6 and MCL-1 during *M. avium paratuberculosis* infection (37). Notch-1 signaling was shown to modulate macrophage polarization and immune defense against during infection, but the molecular mechanisms were not defined; however, the molecular mechanisms utilized for TRP120 Notch activation have not been previously studied (6). In this study, we investigated the molecular interactions involved in TRP120 Notch activation and have defined a TRP120 Notch SLiM mimetic responsible for Notch activation.

Molecular mimicry has been well-established as an evolutionary survival strategy utilized by pathogens to disrupt or co-opt host function as a protective mechanism to avoid elimination by the host immune system (30, 32–36). More specifically, SLiMs are a distinct, intrinsically disordered class of protein interaction motifs that have been shown to evolve *de novo* for promiscuous binding to various partners and have been documented as a host hijacking mechanism for pathogens (26, 29, 30). Although SLiM mimicry has been established as a mechanism utilized by pathogens to repurpose host cell functions for survival, a Notch ligand mimic has never been defined.

TRP120 contains four intrinsically disordered tandem repeat (TR) domains that have been previously described as important for TRP120’s moonlighting capabilities (9, 12). Within these intrinsically disordered domains are various SLiMs responsible for TRP120 multi-functionality. We have recently defined a novel TRP120 repetitive SLiM that activates Wnt signaling to promote *E. chaffeensis* infection (3). In the current study, we have also determined TRP120-TR as the domain also responsible for Notch activation. Sequence homology studies and Information Spectrum Method (ISM) have shown sequence similarity and similar biological function between TRP120 and endogenous Notch ligands. ISM is a virtual spectroscopy method utilized to predict if proteins share a similar biological function based on the electron-ion interaction potential of amino acids, and only requires the nucleotide sequence of each protein. It was recently used to determine prediction of potential receptor, natural reservoir, tropism and therapeutic/vaccine target of SARS-CoV-2 (38). Our results demonstrate a shared sequence similarity and biological function with both canonical and non-canonical Notch ligands that occurs within the tandem repeat domain of TRP120 (TRP120-TR). Both sequence homology and ISM studies identified specific tandem repeat sequences that are functionally associated with endogenous Notch ligands and range between 20-35 amino acids in size. This data suggested that intrinsically disordered regions found within the TRP120-TR domain are responsible for Notch ligand mimic function and direct effector-host protein interaction with the Notch receptor.

Notch ligand binding occurs specifically with EGFs 11-13 within the LBR of the Notch receptor (39, 40). Canonical Notch ligands are known to contain a DSL domain that is important for Notch binding and activation, but a conserved activation motif has not been defined. Colocalization of TRP120 with Notch-1 was previously shown to occur during *E. chaffeensis* infection (6); however, a direct interaction was not previously shown using yeast-two hybrid (8), possibly due to limitations of this technique with protein interactions involving membrane proteins (6, 41). Using pull down, SPR and protein-coated fluorescent microsphere approaches, we further studied TRP120-Notch-1 interaction and found direct binding occurs through TRP120-TR at a Notch-1 LBR (EGFs 1-15). TRP120-TR and Notch-1 LBR interaction occurred at an affinity of 120 ± 2.0 nM, indicating a strong protein-protein interaction. Numerous structural studies of interactions of Notch with endogenous ligands have shown low affinity interactions between Notch Jag or DLL ECDs (42–44). One study demonstrated weak affinities between Notch-1 with an engineered high affinity Jag-1 variant (K_D_ = 5.4 μM) and DLL4 (12.8 μM) (39). The higher binding affinity of TRP120-TR in comparison to canonical Notch ligands suggests that the four tandemly repeated motifs folds in a structure that potentiates binding between TRP120 and Notch-1. In addition, stimulating THP-1 cells and primary monocytes with TRP120-TR resulted in NICD nuclear translocation, indicating that TRP120-TR is the TRP120 domain responsible for Notch activation. Interestingly, TRP120-Fzd5 interaction also occurred through the tandem repeat domain and supports our current findings that TRP120-host protein interactions occur within regions of the tandem repeat domain, likely due to its disordered nature (3).

Secreted and membrane-bound proteins have been shown to activate Notch signaling. These non-canonical Notch ligands lack the DSL domain but still have the ability to modify Notch signaling. Some of the non-canonical Notch proteins contain EGF-like domains; however, others share very little sequence similarity to endogenous Notch ligands (23, 45). TSP2 is a secreted mammalian protein containing EGF-like domains. TSP2 was found to potentiate Notch signaling by direct Notch-3/Jagged1 binding (46). Furthermore, TSP2 binds directly to purified Notch-3 protein containing EGF-like domains 1–11, suggesting a direct interaction. Non-canonical Notch ligand TSP2 was found to share significant sequence homology within the TRP120-TR sequence. Homologous regions included the identified TRP120-TR Notch SLiM mimetic. Although TSP2 has been identified as a secreted, non-canonical Notch ligand, there has been no activating motif identified to date. F3/contactin1, another identified secreted non-canonical Notch ligand, does not contain DSL or EGF-like domains; however, it activates the Notch signaling pathway through the Notch-1 receptor (47). TRP120 was found to share biological function with F3/contactin1 by ISM. F3/contactin1 has been demonstrated to bind to Notch-1 at two different locations within the NECD and activates Notch signaling when presented as purified soluble protein (47). Therefore, Notch activation by secreted, non-canonical Notch ligands has been demonstrated; however, more insight into the molecular details of those interactions needs to be elucidated. This study provides new insight regarding non-canonical Notch ligand activation of the Notch signaling pathway.

SLiMs have been identified in secreted effector proteins of intracellular bacterial pathogens, including *Ehrlichia, Anaplasma phagocytophilum* (48), *Legionella pneumophila* (49–51) and *Mycobacterium tuberculosis* (52). This investigation identifies a novel Notch SLiM (11 aa) that can activate Notch signaling as a soluble ligand. Complete NICD nuclear translocation was previously shown to occur at 2 h post-infection (6), indicating that NICD nuclear translocation during *E. chaffeensis* infection is a result of TRP120-TR Notch ligand SLiM mimetic interaction with the Notch-1 receptor. In addition, Notch signaling pathway genes were upregulated at 24 h in TRP120-TR Notch mimetic SLiM-treated THP-1 cells. These data are consistent with our previous findings where we detected upregulation of Notch signaling pathway components and target genes during *E. chaffeensis* infection at 12, 24, 48, and 72 h.p.i., with maximum changes in Notch gene expression occurring at 24 h.p.i (6). Furthermore, during *E. chaffeensis* infection, TRP120 mediated ubiquitination and proteasomal degradation of Notch negative regulator, FBW7 begins at 24 h.p.i. and gradually decreases during late stages of infection (2). Both TRP120 and FBW7 are localized to the nucleus beginning at 24 h.p.i., suggesting that TRP120-degradadtion of FBW7 assists in upregulation of Notch downstream targets at this timepoint (2).

Interestingly, both the TRP120 Notch memetic IDD (TRP120-N1-P3) and SLiM (TRP120-N1-P6) resulted in concentration-dependent upregulation of Notch downstream targets. Similar to our findings, studies have shown that the Notch pathway can induce heterogenous phenotypic responses in a Notch ligand or NICD dose dependent manner. Klein et al. demonstrated that high levels of Notch ligands can induce a ligand inhibitory effect, while lower levels of Notch ligand activate Notch signaling activity (53). Similarly, Semenova D et al. has shown that NICD and Jag1 transduction increases osteogenic differentiation in a dose-dependent manner; however high dosage of NICD and Jag1 decreases osteogenic differentiation efficiency (54). Furthermore, Gomez-Lamarca et al. has shown that NICD dosage can influence CSL-DNA binding kinetics, NICD dimerization, and chromatin opening to strengthen transcriptional activation (55). Therefore, an increase in Notch ligand-receptor interaction may lead to increased NICD release and Notch signaling strength.

Alanine mutagenesis demonstrated the entire 11-aa TRP120-TR Notch ligand SLiM mimetic is required for Notch activation. Importantly, SLiMs are known to have low-affinity, transient protein-protein interactions within the low-micromolar range (26). In this case, the repeated TRP120-TR Notch ligand SLiM mimetic motif may cause TRP120 to fold in a tertiary structure upon binding to the Notch-1 receptor that stabilizes the TRP120-Notch-1 interaction. Based on this data, *E. chaffeensis* TRP120 could be used as a model to study SLiMs within intrinsically disordered effector proteins that are utilized for host exploitation by other intracellular bacterial pathogens.

To demonstrate that TRP120-TR Notch ligand SLiM mimetic motif is solely responsible for *E. chaffeensis* activation of Notch, we generated an antibody against the mimetic epitope to block *E. chaffeensis* TRP120-Notch-1 binding. Our results demonstrated antibody blockade of Notch activation by *E. chaffeensis*, rTRP120 and the TRP120-TR Notch ligand SLiM peptide. This data strongly supports the conclusion that the TRP120-TR Notch ligand SLiM mimetic is responsible for *E. chaffeensis* Notch activation and may provide a new *E. chaffeensis* therapeutic target. Hence, this study serves to provide insight into the molecular details of how Notch signaling is modulated during *E. chaffeensis* infection and may serve as a model for other pathogens.

Further outstanding questions regarding regulation of the Notch signaling pathway during *E. chaffeensis* remain. We have recently demonstrated maintenance of Notch activation is linked to TRP120-mediated ubiquitination and proteasomal degradation of tumor suppressor FBW7, a Notch negative regulator (2). However, other potential Notch regulators may serve as a target for TRP120-medidated ubiquitination for constitutive Notch activation during infection. Suppressor of Deltex [Su(dx)] is an E3 ubiquitin ligase that serves as another negative regulator of Notch signaling by degrading Deltex, a positive regulator of Notch signaling (56). Su(dx) may serve as another target of TRP120-mediated ubiquitination to maintain Notch activity during *E. chaffeensis* infection. Furthermore, how secreted non-canonical Notch ligands are able to cause separation between the NICD and NECD remains unknown. TRP120 causes Notch activation, resulting in upregulation of Notch downstream targets; however, the mechanism of how the S2 exposure for ADAM cleavage is not understood. Future crystallography studies on TRP120 and Notch-1 interaction may provide more insight into these structural details required for TRP120-N1-P6 SLiM Notch activation (57, 58).

In conclusion, we have demonstrated *E. chaffeensis* Notch activation is initiated by a TRP120 Notch SLiM mimetic. Our findings have identified a pathogen protein host mimic to repurpose the evolutionarily conserved Notch signaling pathway for intracellular survival. This study gives more insight into how obligate intracellular pathogens, with small genomes have evolved host mimicry modules *de novo* to exploit conserved signaling pathways to suppress innate defenses to promote infection.

## Materials and Methods

### Cell culture and cultivation of *E. chaffeensis*

Human monocytic leukemia cells (THP-1; ATCC TIB-202) were propagated in RPMI media (ATCC) containing 2 mM L-glutamine, 10 mM HEPES, 1 mM sodium pyruvate, 4500 mg/L glucose,1500 mg/L sodium bicarbonate, supplemented with 10% fetal bovine serum (FBS; Invitrogen) at 37°C in 5% CO_2_ atmosphere. *E. chaffeensis* (Arkansas strain) was cultivated in THP-1 cells. Host cell-free *E. chaffeensis* was prepared by rupturing infected THP-1 cells or primary human monocytes with sterile glass beads (1 mm) by vortexing. Infected THP-1 cells were harvested and pelleted by centrifugation at 500 × *g* for 5 min. The pellet was resuspended in sterile phosphate-buffered saline (PBS) in a 50-ml tube containing glass beads and vortexed at moderate speed for 1 min. The cell debris was pelleted at 1,500 × *g* for 10 min, and the supernatant was further pelleted by high-speed centrifugation at 12,000 × *g* for 10 min, 4°C. The purified ehrlichiae were resuspended in fresh RPMI media and utilized as needed.

### Human PBMC and primary monocyte isolation

Primary human monocytes were isolated from 125ml of human blood obtained from Gulf Coast Regional Blood Center (Houston, TX). Blood was diluted in RMPI media and separated by density gradient separation on Ficoll at 2000rpm for 20 minutes. The plasma was removed from the separated sample and the buffy coat was collected. Buffy coat was diluted with DPBS containing 2% FBS and 1mM EDTA and centrifuged at 1500rpm for 15 minutes. Supernatant was removed and all cells were combined and mixed carefully. Combined cells were then centrifuged at 1500rpm for 10 minutes and supernatant was removed. Cells were resuspended into 1mL of DPBS containing 2% FBS and 1mM EDTA. Cells were then diluted to 5 × 10^7^/mL concentration, and monocytes were separated by the EasySep Human Monocyte Enrichment Kit w/o CD16 depletion (Stemcell #19058) according to the manufacturers protocol. Primary human monocytes were then cultured in RPMI media (ATCC) containing 2 mM L-glutamine, 10 mM HEPES, 1 mM sodium pyruvate, 4500 mg/L glucose,1500 mg/L sodium bicarbonate, supplemented with 10% fetal bovine serum (FBS; Invitrogen) at 37°C in 5% CO_2_ atmosphere.

### Antibodies and Reagents

Primary antibodies used in this study for immunofluorescence microscopy, Western blot analysis, and pull-down assays include monoclonal rabbit α-Notch1 (3608S; Cell Signaling Technology, Danvers MA), polyclonal rabbit α-Notch1, intracellular (07-1231; Millipose Sigma, Billerica, MA), rabbit α-TRP120-I1 (59) polyclonal rabbit α-TRX (T0803; Sigma-Aldrich, Saint Louis, MO). Polyclonal rabbit anti-TRP120 antiserum was commercially generated against a TRP120 epitope inclusive of aa. 290-301 (GenScript, Piscataway, NJ). Synthetic peptides used in this study were commercially generated (Genscript, Piscataway, NJ).

### Sequence Homology

Genome and transcriptome sequences encoding *E. chaffeensis* TRP120 and *Homo sapiens* Notch ligand proteins were recovered using BLAST searches with the online version at the NCBI website. Sequences were submitted to NCBI Protein BLAST and ClustalW2 sequence databases for sequence alignment.

### Informational Spectrum Method (ISM)

ISM analyzes the primary structure of proteins by assigning a physical parameter which is relevant for the protein’s biological function (38, 60). Each amino acid in TRP120 and Notch ligand sequences was given a value corresponding to its electron-ion interaction potential (EIIP), which determines the long-range properties of biological molecules. The value of the amino acids within the protein were Fourier transformed to provide a Fourier spectrum that is representative of the protein, resulting in a series of frequencies and amplitudes. The frequencies correspond to a physico-chemical property involved in the biological activity of the protein. Comparison of proteins is performed by cross-spectra analysis. Proteins with similar spectra were predicted to have a similar biological function. Inverse Fourier Transform was performed to identify the sequence responsible for obtained signals at a given frequency.

### Transfection

HeLa cells (1 × 10^6^) were seeded in a 60 mm culture dish 24 h prior to transfection. AcGFP-TRP120 or AcGFP-control plasmids were added to OptiMem and Lipofectamine 2000 mixture and incubated for 20 min at 37°C. Lipofectamine/plasmid mixtures were added to HeLa cells and incubated for 4 h at 37°C. Media was aspirated 4 h post-transfection and fresh media was added to each plate and incubated for 24 h.

### Pull Down Assay

Recombinant His-tagged TRP120 (10 μg) and Notch-1 (10 μg) (Sino Biological) were incubated with Ni-NTA beads alone, or in combination, for 4 h at 4°C. Supernatants were collected and the Ni-NTA beads were washed 5X with 10 mM imidazole wash buffer. Proteins were eluted off with 200 mM imidazole elution buffer and binding determined by Western blot analysis.

### Immunofluorescent Confocal Microscopy

THP-1 cells (2 ×10^6^) were treated with full length or truncated constructs (-TR or -C terminus) of recombinant TRP120, or TRP120 peptides for 2 h at 37°C. Cells were collected and fixed using 4% formaldehyde, washed with 1X PBS and permeabilized and blocked in 0.5% Triton X-100 and 2% BSA in PBS for 30 min. Cells were washed with PBS and probed with polyclonal rabbit α-Notch-1, intracellular (1:100) (Millipore Sigma, MA) or monoclonal rabbit α-Notch1 (3608S; Cell Signaling Technology, Danvers MA) for 1 h at room temperature. Cells were washed with PBS and probed with Alexa Fluor 568 rabbit anti-goat IgG (H+L) for 30 min at room temperature, washed and then mounted with ProLong Gold antifade reagent with DAPI (Molecular Probes, OR). Slides were imaged on a Zeiss LSM 880 confocal laser scanning microscopy. Pearson’s correlation coefficient and Mander’s correlation coefficient was generated by ImageJ software to quantify the degree of colocalization between fluorophores.

### Protein-coated fluorescent microsphere assay

TRP120 and TRX recombinant proteins were desalted using Zeba spin desalting columns (Thermo Fisher Scientific, MA) as indicated by the manufacturer protocol. Protein abundance of desalted recombinant protein was assessed by bicinchoninic acid assay (BCA assay). One-micrometer, yellow-green (505/515), sulfate FluoSpheres (Life Technologies, CA) were first equilibrated with 40μM of MES buffer followed by incubation with 10μg of desalted TRP120 or TRX recombinant protein in 40μM MES (2-(N-morpholino) ethanesulfonic acid) buffer for 2 h at room temperature on a rotor. TRP120 or TRX coated FluoSpheres were washed twice with 40μM MES buffer at 12,000 × g for 5 mins and then resuspended in RPMI media. To determine TRP120 or TRX protein coating of FluoSpheres, dot blotting of FluoSpheres samples was performed after protein coating using α-TRX or α-TRP120 antibodies. 8 × 10^5^ THP-1 cells/well were plated in a 96-well round bottom plate, and the TRP120 or TRX coated FluoSpheres were added to each well at approximately 5 beads/cell. The cell and protein-coated FluoSpheres were incubated between 5-60 mins at 37°C with 5% CO^2^, collected and unbound beads were washed twice with 1 X PBS, followed by fixation by cytospin for 15 mins. Cell samples were then processed for analysis by immunofluorescent confocal microscopy, as previously mentioned. FluoSpheres are light-sensitive, therefore all steps were performed in the dark.

### Quantitative Real-time PCR

The human Notch signaling targets PCR array profiles the expression of 84 Notch pathway-focused genes to analyze Notch pathway status. PCR arrays were performed according to the PCR array handbook from the manufacturer. Briefly, uninfected and *E. chaffeensis*-infected or Notch mimetic peptide-treated THP-1 cells were collected at 24 and 48 h intervals and RNA purification with minor modifications, cDNA synthesis and real-time PCR were performed as previously described (3).

### Western Blot Analysis

Cells were lysed in RIPA lysis buffer (0.5M Tris-HCl, pH 7.4, 1.5M NaCl, 2.5% deoxycholic acid, 10% NP-40, 10mM EDTA) containing protease inhibitor cocktail for 30 min at 4°C. Lysates were then cleared by centrifugation and protein abundance assessed by bicinchoninic acid assay (BCA assay). Samples were added to Laemelli buffer then boiled for 5 min. Lysates were then subjected to SDS-PAGE followed by transfer to nitrocellulose membrane. Membranes were blocked for 1 h in 5% nonfat milk diluted in TBST and then exposed to α-TRP120, α-TRX or α-Notch-1 primary antibodies overnight. Membranes were washed three times in Tris-buffered saline containing 1% Triton (TBST) for 30 min followed by 1 h incubation with horseradish peroxidase-conjugated anti-rabbit and anti-mouse secondary antibodies (SeraCare, Milford, MA) (diluted 1:10,000 in 5% nonfat milk in TBST). Proteins were visualized with ECL via Chemi-doc2 and densitometry was measured with VisionWorks Image Acquisition and Analysis Software.

### Surface Plasmon Resonance

SPR was performed using a BIAcore T100 instrument with nitrilotriacetic acid (NTA) sensor chip. Purified polyhistidine-tagged, full-length, rTRP120-TR, rTRX and human rNotch-1 Fc Chimera Protein, CF (R&D Systems, MN) were dialyzed in running buffer (100 mM sodium phosphate [pH 7.4], 400 mM NaCl, 40 μM EDTA, 0.005% [vol/vol]). Briefly, each cycle of running started with charging the NTA chip with 500 μM of NiCl^2^. Subsequently, purified polyhistidine-tagged, full-length, truncated rTRP120 proteins, or rTRX (0.1 μM) were immobilized on the NTA sensor as ligand on flow cell 2. Immobilization was carried out at 25°C at a constant flow rate of 30 μl/min for 100s. Varying concentrations of Notch1-NECD constructs (0-800 nM) were injected over sensor surfaces as analyte with duplicates along with several blanks of running buffer. Injections of analyte were carried out at a flow rate of 30 μl/min with contact time of 360 s and a dissociation time of 300 s. Finally, the NTA surface was regenerated by using 350 mM EDTA. Readout included a sensogram plot of response against time, showing the progress of the interaction. Curve fittings were done with the 1:1 Langmuir binding model with all fitting quality critique requirements met. The binding affinity (K_D_) was determined for all interactions by extracting the association rate constant and dissociation rate constant from the sensorgram curve (K_D_ = Kd/Ka) using the BIAevaluation package software.

### TRP120 Antibody Inhibition of *E. chaffeensis* Notch activation

Host cell-free *E. chaffeensis* was pre-treated with 5-10 μg/ml of polyclonal rabbit anti-TRP120 antibody generated against the TRP120 Notch mimetic SLiM (aa. 284-301), or purified IgG antibody. The cell-free *E. chaffeensis/antibody* mixture was then added to THP-1 cells (5 × 10^5^) in a 12-well plate for 2 h. Samples were collected, washed with PBS and prepared for IFA.

### TRP120 Protein Expression and Purification

Full length or truncated constructs of rTRP120, or rTRX control were expressed in a pBAD expression vector, which has been previously optimized by our laboratory (59, 61, 62). Recombinant TRP120 full length, truncated constructs, and rTRX were purified via nickel-nitrilotriacetic acid (Ni-NTA) purification system. All recombinant proteins were dialyzed via PBS and tested for bacterial endotoxins using the Limulus Amebocyte Lysate (LAL) test.

### Statistical Analysis

All data are represented as the means ± standard deviation (SD) of data obtained from at least three independent experiments done with triplicate biological replicates, unless otherwise indicated. Analyses were performed using a two-way ANOVA or two-tailed Student’s *t*-test (GraphPad Prism 6 software, La Jolla, CA). A P-value of <0.05 was considered statistically significant.

## Acknowledgments

We thank the UTMB Solution Biophysics Laboratory and the Optical Microscopy Core for assistance with confocal microscopy. This work was supported by the National Institute of Allergy and Infectious Diseases grants AI158422, AI149136 and AI126144 to J.W.M., a UTMB McLaughlin Endowment Predoctoral Fellowship to L.L.P., an NIH 1F31AI152424 fellowship to L.L.P., and a T32AI007526-20 biodefense training fellowship to C.D.B.

---

**Fig. S1.**
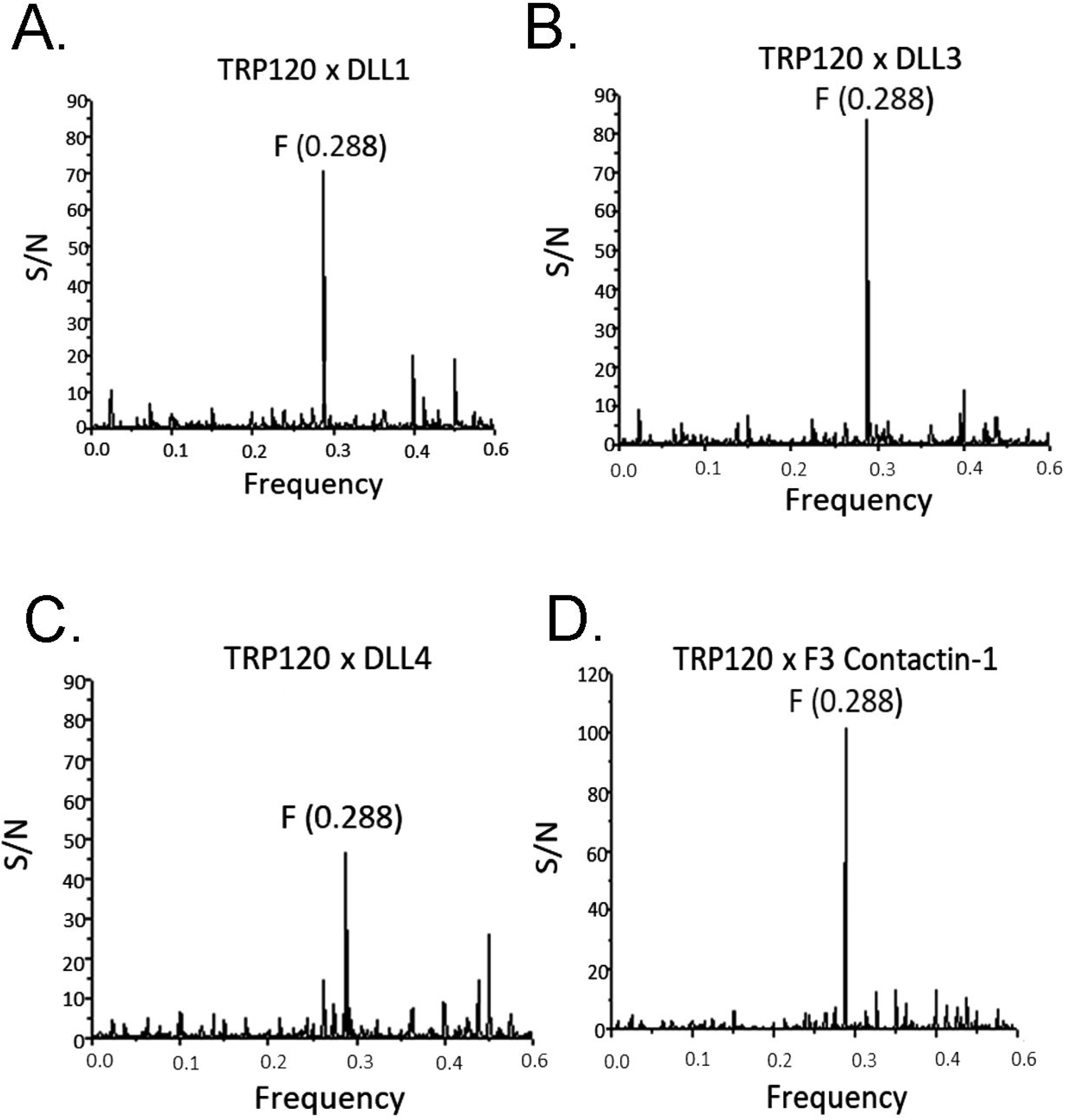
TRP120 shares biological function with canonical and noncanonical Notch ligands. Informational Spectrum Method (ISM) was used to predict if TRP120 shared similar biological function with endogenous canonical and noncanonical Notch ligands. The primary sequence of TRP120 and endogenous Notch ligands were converted into a numerical sequence using each amino acids electron ion interaction potential (EIIP). Numerical sequences were converted into a spectrum using Fourier transform. To determine if proteins shared a similar biological function and cross spectra analysis was performed with TRP120 and Notch ligands individually. Similar biological function is denoted by a peak at a frequency of F(0.288). TRP120 was predicted to share a similar biological function as canonical Notch ligands (A) DLL1, (B) DLL3 and (C) DLL4 and (D) noncanonical Notch ligand, F3 Contactin-1.

**Fig. S2.**
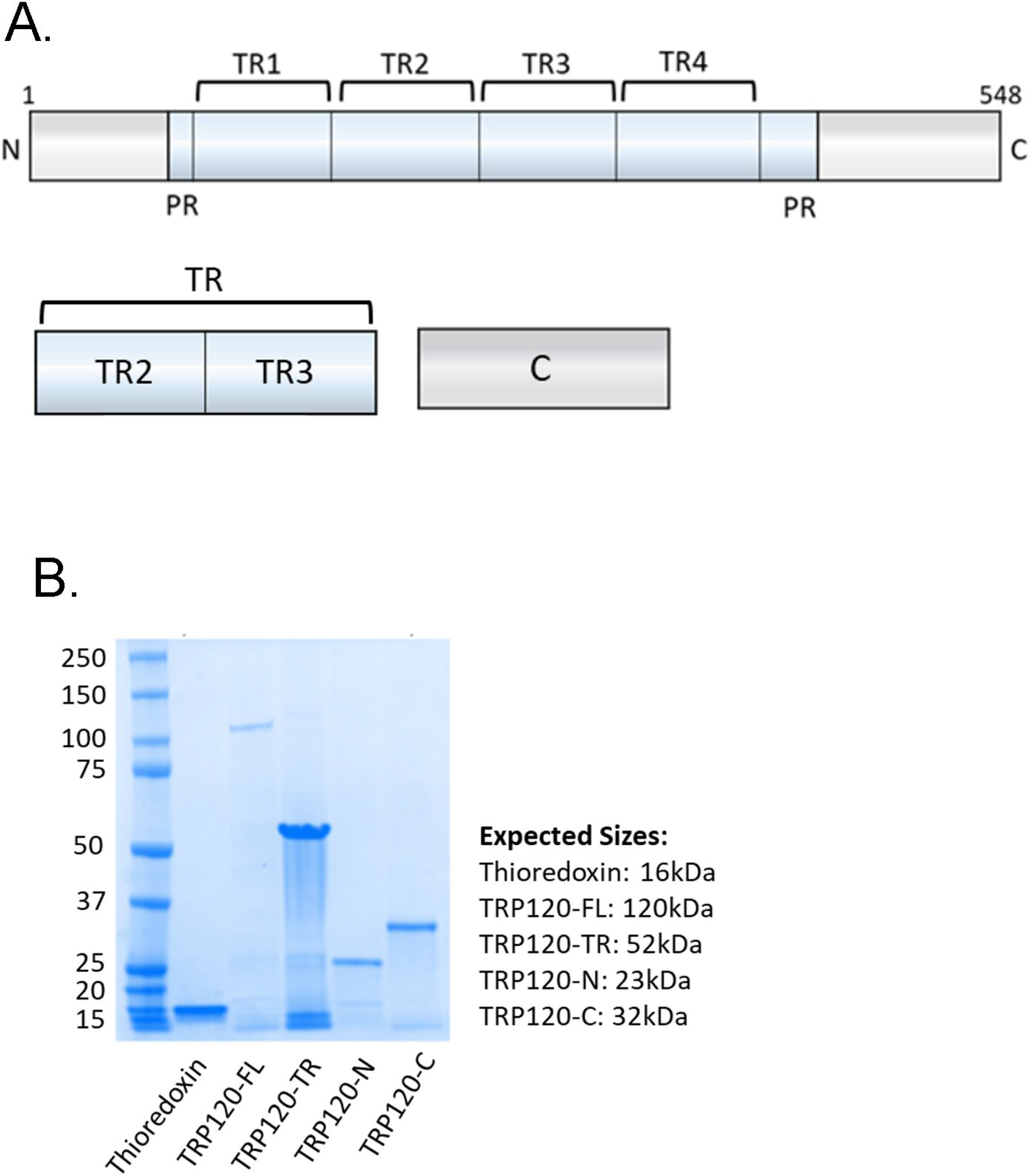
Purification of recombinant TRP120 proteins. (A) Schematic of TRP120-FL, -TR and −C-terminus recombinant proteins. TRP120-TR is expressed and purified as two tandem repeat domains. (B) Coomassie Blue stained gel displaying expression of purified TRP120-FL, - TR, -N, −C-terminus and TRX recombinant proteins. All listed recombinant proteins were expressed in a pBAD vector containing a His-tag.

**Fig. S3.**
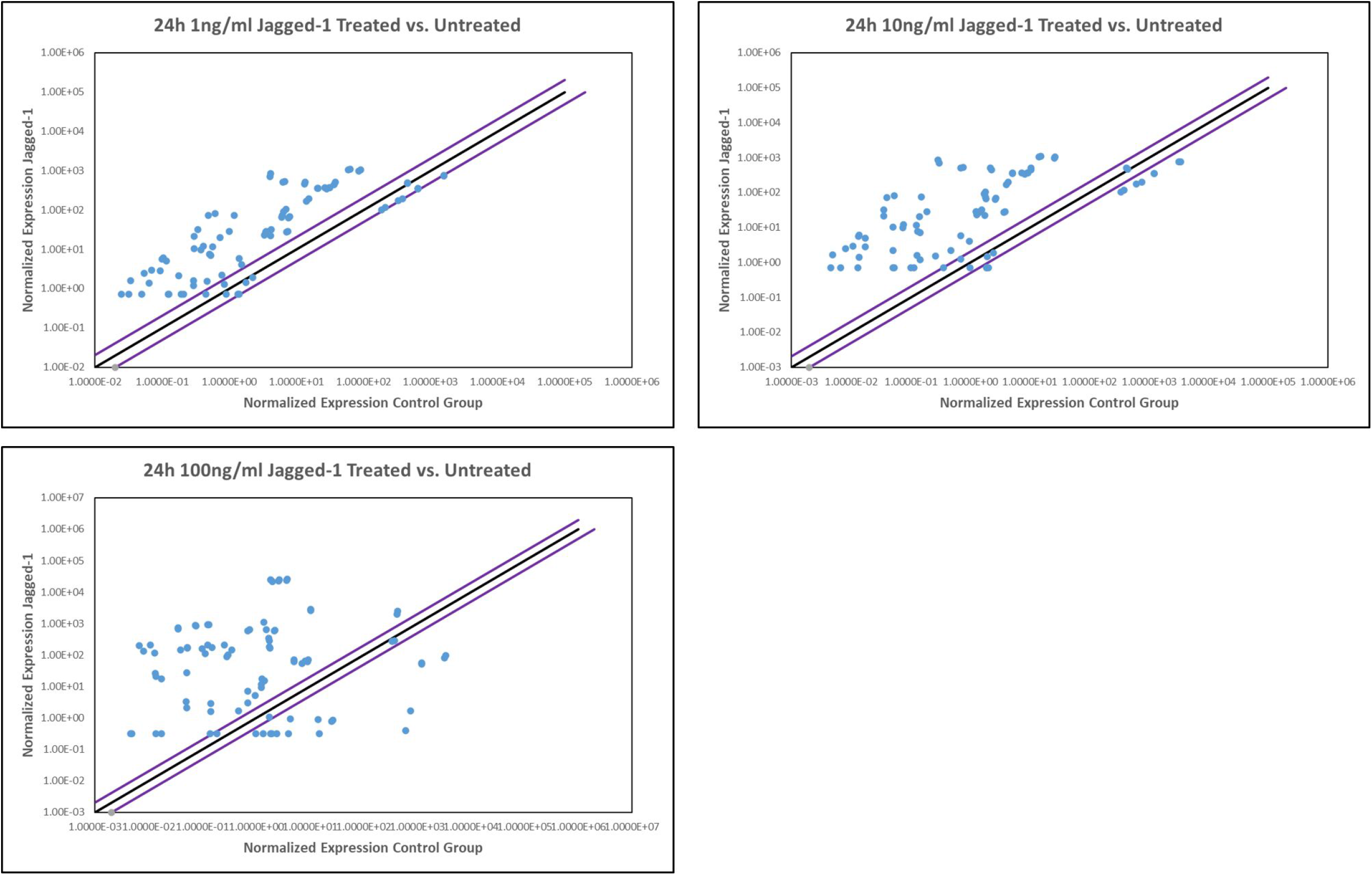
Jagged-1 activates Notch gene expression in a concentration-dependent manner. Scatter plots of expression array analysis of 84 Notch signaling pathway genes to determine Notch gene expression with 1 ng/ml (top left). 10 ng/ml (top right) 100 ng/ml (bottom left) of recombinant Jagged-1 at 24 p.t. Purple lines denote a 2-fold up or down regulation in comparison to control, and the black lines denotes no change.

